# Manipulating phloem transport affects wood formation but not nonstructural carbon concentrations in an evergreen conifer

**DOI:** 10.1101/2020.09.25.313569

**Authors:** Tim Rademacher, Patrick Fonti, James M. LeMoine, Marina V. Fonti, David Basler, Yizhao Chen, Andrew D. Friend, Bijan Seyednasrollah, Annemarie H. Eckes-Shephard, Andrew D. Richardson

## Abstract

Wood formation is a crucial process for carbon sequestration, yet how variations in carbon supply affect wood formation and carbon dynamics in trees more generally remains poorly understood.

To better understand the role of carbon supply in wood formation, we restricted phloem transport using girdling and compression around the stem of mature white pines and monitored the effects on local wood formation and stem CO_2_ efflux, as well as nonstructural carbon concentrations in needles, stems, and roots.

Growth and stem CO_2_ efflux varied with location relative to treatment (i.e., above or below on the stem). We observed up to a two-fold difference in the number of tracheids formed above versus below the manipulations over the remaining growing season. In contrast, the treatments did not affect mean cell size noticeably and mean cell-wall area decreased only slightly below them. Surprisingly, nonstructural carbon pools and concentrations in the xylem, needles, and roots remained largely unchanged, although starch reserves declined and increased marginally below and above the girdle, respectively.

Our results suggest that phloem transport strongly affects cell proliferation and respiration in the cambial zone of mature white pine, but has little impact on nonstructural carbon concentrations. These findings contribute to our understanding of how wood formation is controlled.

**Highlight:** Restrictions in phloem transport designed to affect carbon supply, lead to changes in wood formation and stem respiration of mature white pines without substantially changing local nonstructural carbon concentrations.

## 1. Introduction

Forests sequester about 14% of anthropogenic carbon emissions each year (Pan *et al*., 2011) and wood formation is a pivotal process sequestering this carbon for extended periods. Since wood can persist for millennia and is composed of roughly 50% carbon (Lamlom and Savidge, 2003), allocation to wood is a salient carbon sink. Understanding the mechanisms of wood formation is therefore crucial to assess land carbon sequestration in a rapidly changing world (Hartmann *et al*., 2020) and for accurate predictions of the land carbon cycle (Friend *et al*., 2019).

Many studies have focused on assessing the “fate” of assimilated carbon in land plants (Atkin, 2015) with the expectation of a direct link between carbon assimilation and integration into wood structure. Experiments manipulating carbon assimilation by artificially increasing atmospheric CO_2_ (Norby and Zak, 2011), defoliating (Hoch, 2005) or shading trees (O’Brien *et al*., 2014) have often been undertaken to describe and quantify the relation between sources and sinks. Growth sinks and carbon status have been shown to be coupled under low atmospheric CO_2_ (Hartmann *et al*., 2013), severe shading (Weber *et al*., 2019), and defoliation (Piper *et al*., 2015; Wiley *et al*., 2017; Fierravanti *et al*., 2019). However, a meta-analysis found that under increased assimilation due to elevated atmospheric CO_2_ plant growth does not increase proportionally to carbon uptake (Ainsworth and Long, 2005), although most included studies used herbaceous plants. Evidence from tree rings of naturally growing trees suggests a declining response of radial growth to increasing CO_2_ with age (Voelker *et al*., 2006) or no response at all (van der Sleen *et al*., 2015), which has also been corroborated in one CO_2_ enrichment study on mature trees (Körner *et al*., 2005). Generally, smaller trees may be more carbon-limited (Körner, 2003; Hayat *et al*., 2017) due to ontological differences in carbon allocation (Hartmann *et al*., 2018), which could explain discrepancies in growth response between smaller and larger trees. A decoupling of carbon assimilation (i.e., photosynthesis) and fixation (e.g., wood formation) has been hypothesised to be the cause of the discrepancy in carbon uptake and growth responses and mediated by energy reserves, primarily nonstructural carbon (Fatichi *et al*., 2014). Presumably, this decoupling is particularly pertinent for larger trees relying on substantial energy reserves.

There is increasing evidence to support the idea that cambial activity is more sensitive to environmental conditions than photosynthesis (Fromm, 2010; Cabon *et al*., 2020; Vieira *et al*., 2020). For example, cambial activity is directly constrained by water status (Steppe *et al*., 2015; Peters *et al*., 2020) and temperature (Begum *et al*., 2018). Thus, environmental sensitivities of the cambium complicate investigations trying to isolate the influence of carbon supply on wood formation due to differences in environmental conditions in space and time. Nonetheless, cell division, enlargement, and cell-wall deposition have been shown to be affected by changes in carbon status due to natural defoliation (Rossi *et al*., 2009; Deslauriers *et al*., 2016; Castagneri *et al*., 2020). Thus, despite these efforts to quantify the relation between assimilation and growth, the debate on whether carbon supply or demand drive allocation to wood formation is not settled yet (Gessler and Grossiord, 2019).

Quantifying the *in situ* response of the constituent processes of wood formation (i.e., cell division, cell enlargement and cell-wall deposition) across a gradient of carbon supply would help to disentangle the processes acting along the long pathways between carbon assimilation in leaves and integration into wood in mature trees. Focusing the attention on the cambium, where carbon is fixed into woody structure, reduces complexity such as potentially confounding whole-tree feedbacks. Phloem transport manipulations can be used to directly manipulate carbon supply to parts of the cambium (Rademacher *et al*., 2019). Locally, there are three main carbon sinks in stems: growth, stem respiration, and nonstructural carbon pools. Radial growth has been repeatedly inferred to be correlated with carbon supply in previous phloem transport manipulations (Wilson, 1968; Maier *et al*., 2010; Maunoury-Danger *et al*., 2010; De Schepper *et al*., 2011). Stem respiration has also been documented to covary with growth along carbon supply gradients (Daudet, 2004; Maunoury-Danger *et al*., 2010), although questions remain whether observed changes in respiration reflect exclusively changes in energy demand due to variations in growth (Rademacher *et al*., 2019). Finally, nonstructural carbon reserves are accumulated and can be remobilised to fuel metabolism in times of scarcity (Carbone *et al*., 2013; Muhr *et al*., 2018). When carbon supply is limited using low atmospheric CO_2_ (Hartmann *et al*., 2015; Huang *et al*., 2019), drought (Hagedorn *et al*., 2016), girdling (Regier *et al*., 2010; Oberhuber *et al*., 2017), or defoliation (Li *et al*., 2002; Wiley *et al*., 2013), observations indicate that allocation to nonstructural carbon reserves is prioritised over growth. On the other hand, little is known about allocation to growth and storage under enhanced carbon supply. At the whole-tree level, we do know that foliage sugar and even more so starch concentrations increase with elevated atmospheric CO_2_ (Ainsworth and Long, 2005, page 2005; Körner *et al*., 2005; Dawes *et al*., 2013; Schmid *et al*.,2017), whereas girdling increased stem starch concentrations above the girdle without affecting sugar or starch concentrations in foliage (Regier *et al*., 2010). Knowledge about whether reserves in local and connected distal tissues are used under carbon deficits and accumulated under carbon surpluses is crucial to understand carbon allocation to wood formation.

In addition to providing local reserves, osmotically active sugars influence turgor pressure (Holtta *et al*., 2010; Guerriero *et al*., 2014). Alterations to osmotically active sugar concentrations have been suggested to be the main causes for changes in growth of girdled trees (De Schepper and Steppe, 2011). Sugars might also be directly involved in signalling for wood formation, as sugar availability controls the expression of cellular synthesis enzymes and thus play a major role in growth (Riou-Khamlichi *et al*., 2000). Seasonal variations of soluble sugar concentration in the cambium have been hypothesised to be a primary control of intra-annual gradients in cell size and cell-wall thickness (Cartenì *et al*., 2018). Finally, sugars are the primary determinant of respiration in growing tissues (Penning De Vries *et al*., 1979; Maier *et al*., 2010). Experiments that directly manipulate the carbon status in the stem and monitor the effects on wood formation, metabolism, and nonstructural carbon have the potential to improve our understanding of the sensitivity of wood formation to carbon supply.

In this study, we aim to investigate controls on structural carbon integration at the sink location in stems of mature trees while monitoring CO_2_ efflux and variation in nonstructural carbon. In particular, we applied phloem girdling and compression to restrict carbon flow along the stem by cutting or exerting theoretically sufficient pressure to collapse phloem tissues around the stem (Henriksson and Rademacher, 2019). These restrictions can generate a large contrast in local carbon supply without affecting light, temperature, water availability, or nutrient availability for the whole tree. To differentiate between carbon supply compensation from local versus distal reserves and assess the role of potential leakage across the compression zone, we added a double compression treatment to isolate a stem section from distal nonstructural carbon reserves (Fig. 1). The resulting carbon-supply gradient is presumed to range from severe carbon limitation below the girdle, over moderate carbon limitation below the compression (due to some leakage), through moderate carbon supply surplus above the compression, to a larger increase of carbon supply above the girdle. As a result of the induced gradient in carbon supply along the stem of mature forest trees, we wanted to answer the overarching question of how ratios of carbon allocation between growth, stem CO_2_ efflux and nonstructural carbon reserves vary across a gradient in carbon supply to cambial regions. In particular, we hypothesised that (H1) radial wood mass growth will covary with inferred carbon supply due to influence of carbon supply on the number of cells, cell sizes and cell-wall areas in newly forming wood; (H2) stem CO_2_ efflux will covary with inferred carbon supply, and finally, (H3) soluble sugar and starch concentrations will also covary with inferred carbon supply.

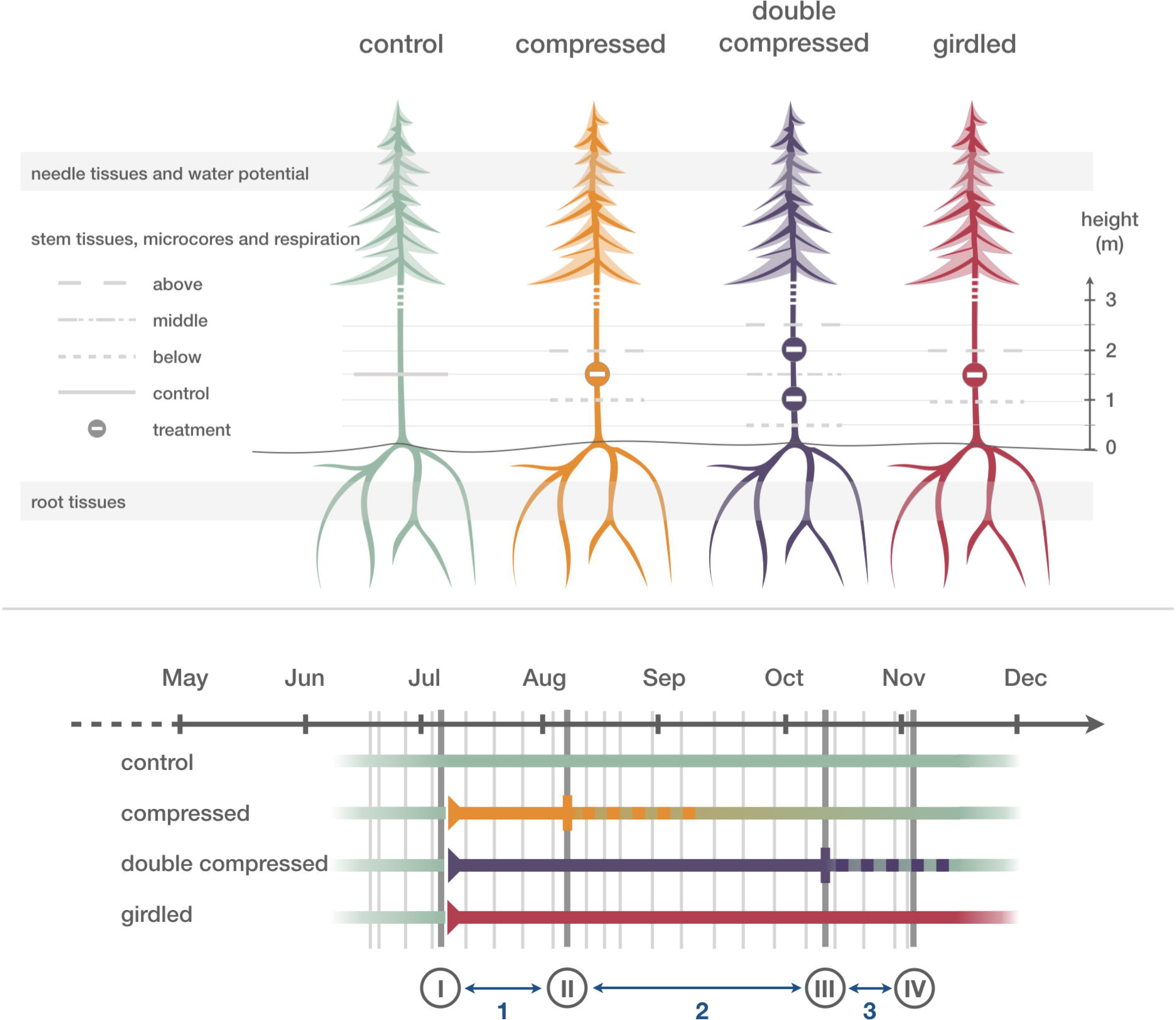

## 2. Materials and Methods

### 2.1 Study site

Harvard Forest is a mesic temperate mixed forest dominated by oak, maple, and pine species located in central Massachusetts, USA. Soils at Harvard Forest are derived from glacial till and characterised as well-draining, slightly acidic sandy loam with an average depth of one meter. Mean annual temperature at the site is 8.0 ± 0.8 °C (μ ± σ) and mean total annual precipitation is 1170 ± 193 mm (μ ± σ), evenly spread across all seasons (Boose and Gould, 2019).

The experiment was conducted on an even-aged cohort of white pines (*Pinus strobus* L.) with homogenous growing conditions. The stand is located at 42°30.8’N and 72°13.1’W, 340 m above sea level, and has been naturally regenerating since a 1990 harvest of a red pine (*Pinus resinosa* Ait.) plantation. Spacing was dense at circa. 1612 trees per hectare. For the experiment, we selected 40, healthy, single-stemmed, co-dominant white pines. At the beginning of the experiment, the selected trees were on average 18 ± 2 years old with a diameter at breast height of 18.7 ± 2.1 cm and a height of 10.9 ± 1.4 m (μ ± σ, see Table 1). The selected trees had grown on average 2.2 ± 1.2 mm y^-1^ in girth and 0.5 ± 0.1 m y^-1^ in height between 2012 and 2017 (μ ± σ).

**Table 1:** Summary statistics for the treatment groups including mean diameter at 1.5 m above the root collar, mean height (*μ* ± 1 *σ* for both), start and end date and duration.

| treatment | n | dbh (cm) | h (m) | start | end | duration (days) |
| --- | --- | --- | --- | --- | --- | --- |
| control | 10 | $18.2 \pm 1.5$ | $10.7 \pm 1.6$ | - | - | - |
| girdled | 10 | $18.9 \pm 2.2$ | $11.1 \pm 1.5$ | 4<sup>th</sup> July | - | - |
| compressed | 10 | $18.2 \pm 2.0$ | $11.4 \pm 0.7$ | 4<sup>th</sup> July | 10<sup>th</sup> August | 36 days |
| double compressed | 10 | $19.7 \pm 2.4$ | $10.5 \pm 1.7$ | 5<sup>th</sup> July | 10<sup>th</sup> October | 95 days |

### 2.2 Experimental setup

One of four treatments - control, single compression, double compression, or girdling - was randomly assigned to each tree, yielding ten trees per treatment. The four treatment groups had similar age, initial diameter at breast height, height, and average radial growth in the previous five years (ANOVA, p = 0.34, 0.46, 0.24, and 0.52, respectively). Dead branches below 3 m were pruned in May 2017 to facilitate access to stems.

The experiment started on the 4^th^ of July 2017. For the control trees, no treatment was applied. For the girdling treatment, we carefully removed a 2.5 cm-wide strip of bark, phloem and cambium around the entire bole at 1.5 m stem height using razor blades (Fig. 1). We did not seal or treat the wounds. For the single compression treatment (hereafter compression treatment), collars were constructed from two ratcheted heavy-duty cargo belts with ratchets diametrically opposite each other as described in (Henriksson and Rademacher, 2019). Compression collars were removed after 36 days (10^th^ August) to allow for potential recovery within the same growing season. For the double compression treatment, we installed identical compression belts at 1.0 m and 2.0 m stem height (Fig. 1) to isolate a stem section with regard to phloem transport to and from the canopy and the roots. Pressure below the belts was measured weekly using piezo-electric pressure sensors (Tactilus Free Form 12mm, Sensor Products Inc., Madison, New Jersey, USA). The belts generated a pressure exceeding 2 MPa around the entire circumference (Fig. S1). We re-tightened the double compression belts after 38 days (13^th^ August) to assure continually sufficient pressures (i.e., > 2MPa) to collapse the phloem, despite loosening over time (Fig. S1a). We finally removed the compression belts after 95 days (10^th^ October). We extended the duration of the double compression versus the single compression to investigate whether recovery was still possible toward the end of the growing season.

### 2.3 Experimental monitoring

All trees were monitored during the experiment to characterise the carbon status of the trees (Fig. 1). Monitoring included a four-time characterisation of tree-ring formation and nonstructural carbon concentrations in needle, stem, and root tissues, and a weekly survey of stem CO_2_ efflux until late November 2017. Two follow-up campaigns of CO_2_ efflux were conducted in 2018 and an additional set of micrcores was collected in 2018. Additionally, tree water status was measured weekly to evaluate potential collateral impacts (see Fig. S2 for details).

Tree-ring growth was characterised from stem microcores collected at heights shown in Fig. 1 with a Trephor (Rossi *et al*., 2006). We collected microcores just before the experiment (3^rd^ July), twice during the experiment (8^th^ August and 10^th^ October), and in late autumn of 2017 and 2018 (3^rd^ November and 1^st^ November, respectively). The microcores were stored in Eppendorf tubes containing a 3:1 solution of ethanol and glacial acetic acid. After 24 hours, the solution was replaced by 95% ethanol. Micro-sections (7 μm-thick cross-sectional cuts) were cut with a rotary microtome (Leica RM2245, Switzerland) from paraffin-embedded samples (Tissue Processor 1020 Leica, Switzerland) and double-stained with astra-blue and safranin. Ring widths were measured using the Wood Image Analysis and Database platform (Rademacher *et al*., in prep.) on microsection images captured using a digital slide-scanner (Zeiss Axio Scan.Z1, Germany) with a resolution of roughly 1.5 pixels per μm. Tracheid anatomical characteristics (e.g., averages of lumen width and cell-wall area over 20 μm-wide tangential bands) were obtained using ROXAS 3.0.285 (von Arx and Carrer, 2014) from the November 2017 microsections.

Stem CO_2_ efflux was measured weekly as an indicator of metabolic activity. For this purpose, chambers (10 cm diameter by 10 cm length PVC pipe) were cut to fit each tree’s curvature at all sampling heights (Fig. 1) and attached two weeks prior to the treatments’ start using silicone adhesive. Starting one week before the beginning of the treatments (29^th^ June), an infrared gas analyser (LI-820, LI-COR, Lincoln, Nebraska, USA) with a circulating pump (12K, Boxer, Ottobeuren, Germany) was attached using a PVC cap with two ports to constantly circulate air through the closed system (Carbone *et al*., 2019). Once the concentration stabilised after closing the lid, the chamber CO_2_ concentration was measured at 1 Hz for at least one minute. Precautions were taken to minimise any effect of diel and environmental influences on treatment differences in CO_2_efflux (Supplementary Information 5). The raw stem CO_2_ efflux and uncertainties were estimated using the RespChamberProc package (http://r-forge.r-project.org/projects/respchamberproc/) as developed by Perez-Priego *et al*. (2015).

Soluble sugar and starch concentrations in coarse roots, stems, and needles were determined from tissue samples collected at the same time as the microcores (Fig. 1). Coarse roots (at least 20 cm below the root collar) and stems were cored using a 5.15 mm increment borer (Haglӧf Company Group, Långsele, Sweden). Foliage samples were collected from a sun-exposed part of the crown with a pole pruner. All sampled tissues were immediately shock-frozen on dry ice in the field and subsequently stored in a freezer (−60°C) until being cut using razor blades and freeze-dried (FreeZone 2.5, Labconco, Kansas City, Missouri, USA and Hybrid Vacuum Pump, Vaccubrand, Wertheim, Germany). Dried samples were ground by machine (mesh 20; Thomas Scientific Wiley Mill, Swedesboro, New Jersey, USA; SPEX SamplePrep 1600, MiniG, Metuchen, New Jersey, USA), although small samples were ground with pestle and mortar to minimise loss of material. An equal mix of first- and second-year needles (more than 100 needles from several branchlets) and the entire root core were each homogenised. For stems, we homogenised the first and second centimetre of the core separately. Only July and November were analysed for the second centimetre. About 40 mg of finely ground and dried powder for all tissue, tree and sampling date combinations was analysed following the protocol by Chow and Landhäusser (2004) as adapted by Furze *et al*. (2019). Colourimetric analysis with phenol-sulphuric acid was read twice at 490 nm for sugar and 525 nm for starch using a spectrophotometer (Thermo Fisher Scientific GENESYS 10S UV-Vis, Waltham, Massachusetts, USA), and calibrated with a 1:1:1 glucose:fructose:galactose (Sigma Chemicals, St. Louis, Missouri, USA) standard curve for sugar and a glucose (Sigma Chemicals, St. Louis, Missouri, USA) standard curve for starch. Each batch of samples included on average 35 samples, at least 10 blanks - both tube and sample blanks - and between 9 and 12 laboratory control standards (red oak stem wood, Harvard Forest, Petersham, Massachusetts, USA; potato starch, Sigma Chemicals, St. Louis, Missouri, USA). We repeated extractions for batches that showed substantial deviations in the laboratory control standards (e.g., starch recovery fraction lower than 85%). The coefficient of variation for laboratory control standards was 0.08 and 0.09 for sugar and starch concentrations in oak wood, respectively, and 0.13 for potato starch. All samples’ absorbance values were converted to concentrations in % dry weight and uncertainties with the self-developed R package NSCprocessR (https://github.com/TTRademacher/NSCprocessR).

To monitor tree water status, we measured pre-dawn needle and branch water potentials once per week per tree from the end of June to the beginning of November. Neither the two compression treatments, nor the girdling, affected tree water status (see Fig. S2 for more details).

### 2.4 Comparing carbon pools and fluxes in a stem section

We scaled structural growth, respiratory losses, and net changes in soluble sugars and starch reserves in the first two centimetres of wood (hereafter stem sugar and starch reserves) to a common unit of grams of carbon in a 10 cm height stem section for each experimental period (sensu Fig. 1) to directly compare the size of these carbon fluxes. For structural carbon increments, we associated the cell-wall area for each 20-*μ*m wide band with a date of formation using the fraction of the ring grown derived from the microsections (i.e., July, August, October). Cell-wall area was then divided by the width of the microcore and multiplied by the circumference and height of the stem section to get the cell-wall volume in the section. Finally, cell-wall volume was multiplied with a cell-wall carbon-density to estimate the mass of carbon fixed in the section (Supplementary Information 4). For losses due to CO_2_ efflux, we averaged CO_2_ efflux rates measured weekly for each period and multiplied them with the surface area of each stem section (Supplementary Information 5). To estimate nonstructural carbon reserves, we multiplied the average soluble sugar and starch concentration in the first and second centimetre of the xylem, which are assumed to be a large and most easily accessible fraction of radial nonstructural carbon reserves (Richardson *et al*., 2015; Furze *et al*., 2020), with the volume of that tissue for each stem section assuming the stem section was perfectly cylindrical. The net change in nonstructural carbon pools for a period is then computed as the difference between each pool’s size at the end and the beginning of each period (Supplementary Information 6).

### 2.5 Statistical analysis

We estimated treatment effects by fitting linear mixed-effects models with the lme4 package (Bates *et al*., 2015) in R v3.6.3 (R Core Team, 2019). A tree identifier was included as a random effect to account for natural between-tree variability, and a fixed effect composed of an interaction between treatment and sampling height (e.g., above, middle and below) was added to substitute for the presumed carbon-supply gradient. When effects over time were estimated, we added date as a categorical fixed effect and to the treatment-sampling height interaction. Models were fitted using restricted maximum likelihood estimation. The strength and importance of estimated effects was judged in comparison with estimated variances. Hereafter, we report estimated effects 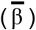, their standard errors 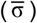, and the *t*-value in the following format: 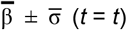. All code and data are publicly available on the Harvard Forest Data Archive (Rademacher and Richardson, 2020).

## 3. Results

### 3.1 Effects on carbon allocation

The treatments had clear effects on growth and CO_2_ efflux, but not on nonstructural carbon pools (Fig. 2). The mass of new wood growth mirrored the presumed carbon-supply gradient. Substantially more carbon (20%, 139%, and 92% respectively) was sequestered in woody tissue above the compression, double compression, and girdle relative to the control from July to November (Fig. 2). Below both compression and double compression, mass growth was not discernibly different from the control from July to November. However, wood formation ceased completely below the girdle after roughly one month after the girdling, approximately halving carbon sequestration in woody structure during the experimental period. CO_2_ efflux was higher above all treatments (46% above compression, 124% above double compression, and 111% above the girdle) between July and November relative to the control. Stem CO_2_ efflux was also substantially reduced below treatments (−45% for compression, −42% for double compression, and −70% for girdling) relative to the control, even when mass growth and the number of forming cells was not noticeably impacted below the compression treatments (Fig. 2, 3). Over the same period, net changes in nonstructural carbon pools in the stems were more than an order of magnitude smaller than the estimated carbon allocation to growth and stem CO_2_ efflux (Fig. 2). A detectable treatment effect on nonstructural carbon concentrations was only apparent in girdled trees.

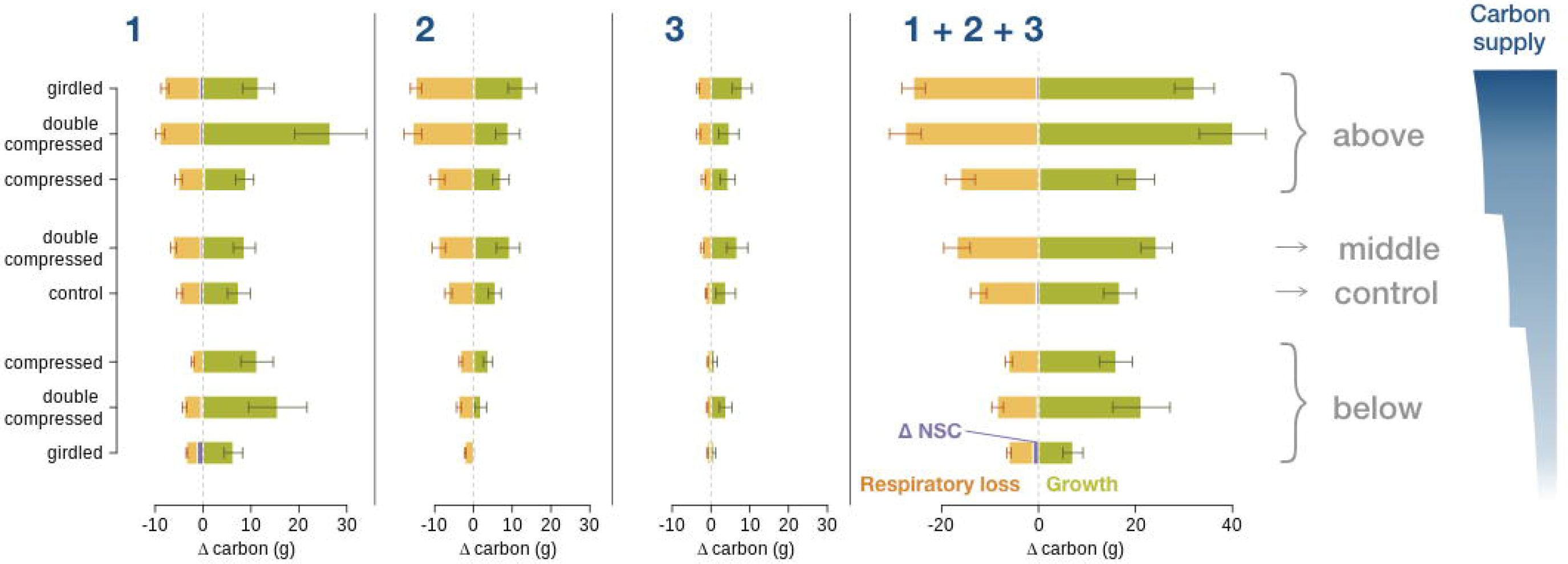

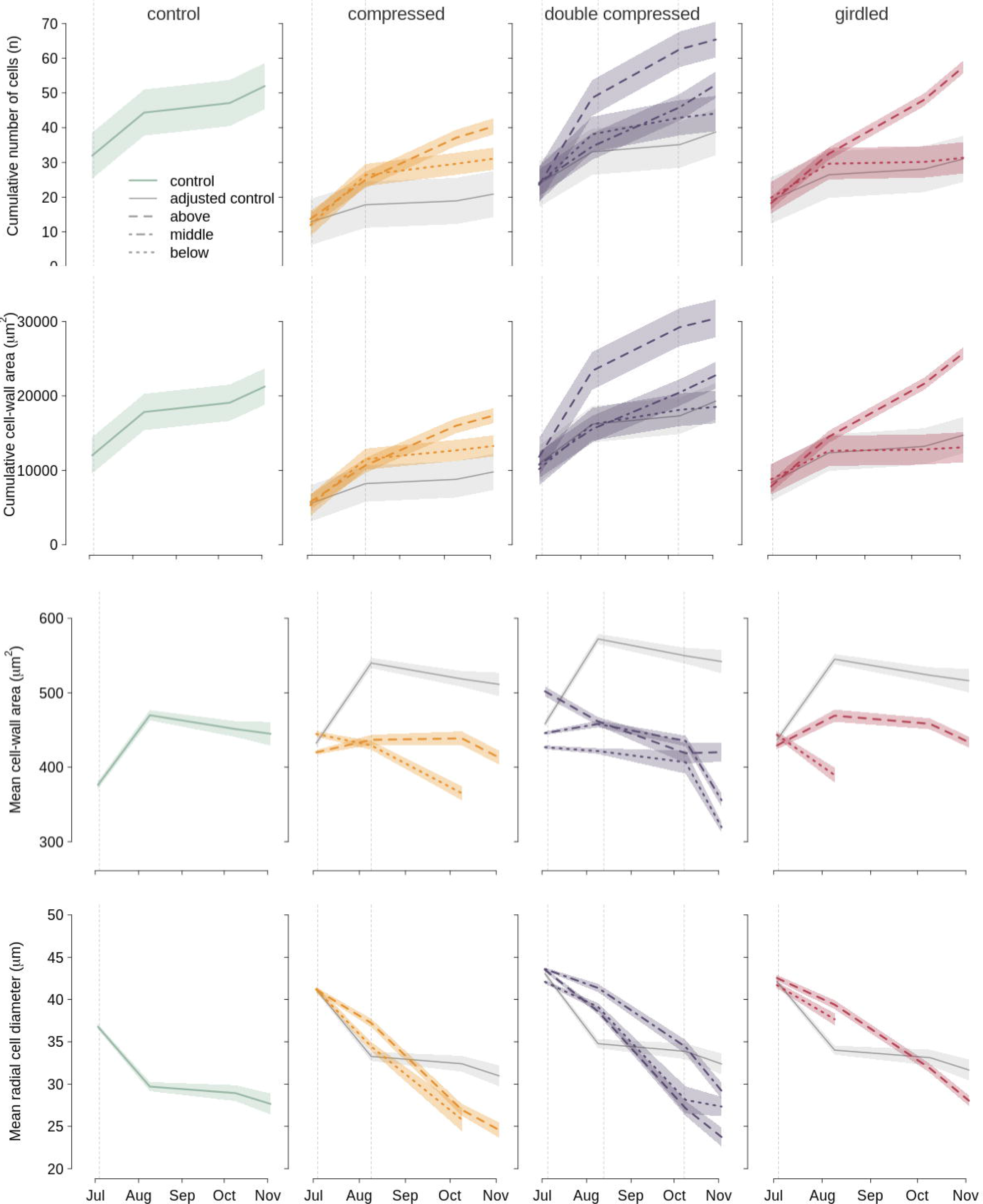

### 3.2 Resulting wood characteristics

About 60.5 ± 3.4% of the final ring width had already formed by the experimental onset (4^th^ July), which coincided with the seasonal peak of wood growth. Control trees showed a typical progression toward smaller tracheids with thicker cell walls over the remainder of the growing season (Fig. 3), resulting in an average of 51 cells per radial file composing a 1.28 mm-wide ring for 2017.

Above the treatments, wood growth was stimulated in the double compression and girdled trees. The final ring was on average 0.73 ± 0.18 (*t* = 5.24), and 0.57 ± 0.18 (*t* = 4.33) mm wider than the control above the double compression and girdle, respectively. However, ring width was not clearly wider above the compression relative to the control with an additional 0.15 ± 0.18 (*t* = 1.90) mm. Differences in ring width could mainly be traced to changes in the number of cells formed after treatment onset, rather than their sizes. All three treatments formed unequivocally fewer cells below the treatment versus above with a difference of 10 ± 6 for compressed, 21 ± 4 for double compressed, and 27 ± 3 for girdled trees. Control and treated trees had similar radial cell diameters (Fig. 3). Mean cell-wall area declined slightly below treatments, which combinated with large differences in the number of cells per radial file above and below treatments caused substantial differences in cumulative cell-wall area, and hence structural biomass.

Below the girdle marginally narrower rings (−220 ± 176 (*t* = −0.18) *μ*m or −17%) with slightly less mean cell-wall area formed compared to the control (Fig. 3). Relative to control, mean cell-wall area was 75 ± 29 (*t* = −2.55) *μ*m^2^ or 18% less below the girdle by August and 47 ± 32 (*t* = −1.49) *μ*m^2^ or 12% less below the compression by October. After August and October, too few cells formed below the girdle and compression to reliably quantify these trends further. In the middle and below the double compression enough cells formed to detect a pronounced decline in mean cell-wall area of −134 ± 32 (*t* = −4.21) *μ*ms^2^ or 35% and −64 ± 31 (*t* = −2.06) *μ*ms^2^ or 17% less, respectively, by November.

Above the girdle, growth resumed in 2018 for 9 of 10 trees at an average of 120 ± 43 % of the standardised ring width of the control group. However, only two trees showed any sign of growth below the girdle in 2018 at 9 and 81% of standardised ring width. For both compression treatments, 19 out of 20 compressed trees grew radially in 2018 at 78 ± 15 and 69 ± 16% of the control group growth above and below the compression and 107 ± 11, 60 ± 11, and 56 ± 20% of the control group above, in the middle, and below the double compression, respectively.

### 3.3 Alterations of stem CO_2_ efflux

CO_2_ efflux of the control group generally declined after a maximum at the start of the experiment (Fig. 4). Losses of carbon due to CO_2_ efflux generally mirrored mass growth in pattern and magnitude across the gradient of carbon supply, but losses due to CO_2_ efflux were more markedly reduced than growth below both compression treatments (Fig. 2). Treatment effects on CO_2_ efflux lagged two to three weeks behind the treatment onset (Fig. 4). Over the entire treatment period the average stimulation of stem CO_2_ efflux above the treatment amounted to 106%, 198%, and 143% of the control for compression, double compression, and girdling, respectively. Below the compression, double compression, and girdle, CO_2_ efflux fell by on average 33%, 49%, and 36% of the control for the period of the treatment. Between the double compression collars, CO_2_ efflux stayed close to the control with an additional 0.53 ± 0.40 (*t* =1.34) *μ*mol m^-2^ s^-1^. By November, both compression treatments’ CO_2_ efflux rates had converged and remained indistinguishable from the control treatment for the following growing seasons (data not shown).

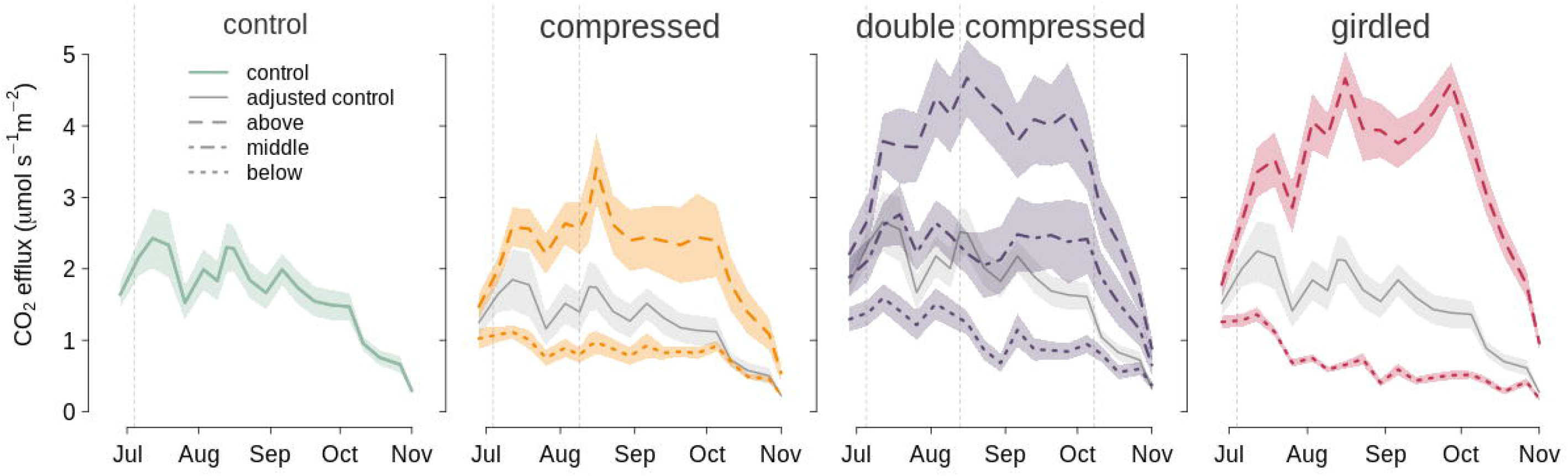

### 3.4 Changes in nonstructural carbon

Nonstructural carbon concentrations, in particular soluble sugars, varied little among treatments and sampling heights. Needle, wood, and root tissues averaged soluble sugar concentrations of 8.33 ± 0.17%, 0.83 ± 0.01%, and 1.47 ± 0.05%, and starch concentrations of 1.32 ± 0.14%, 0.28 ± 0.01%, and 0.31 ± 0.03% across all four measurement dates and trees, respectively. In tissues above all treatments, soluble sugar concentrations mostly followed the control group, but increased slightly in a few tissues (Fig. 5). Most notably, increases of needle soluble sugar concentrations were observed in girdled trees, peaking at an additional 3.30 ± 1.28% (t = 2.58) in November. Needle starch concentrations in girdled trees were also higher in August and October, but had mostly converged (0.44 ± 0.88% (t = 0.49)) with the control by November (Fig. 6). Higher needle soluble sugar concentrations were also apparent for compressed and double compressed trees, albeit with substantially smaller increases than for the girdled trees, culminating in November at 0.56 ± 1.26% (t = 0.44) and 2.38 ± 1.26% (t = 1.88), respectively. Finally, wood sugar concentrations above the treatments increased marginally in the first centimetre by 0.42 ± 0.16% (t = 2.64), 0.35 ± 0.15% (t = 2.22), and 0.23 ± 0.16% (t = 1.44) for compression, double compression and girdling by November, whereas wood starch concentrations remained stable above all treatments but declined below the girdle by 0.32 ± 0.07% (t = −4.48) relative to the control in November. Changes in nonstructural carbon concentrations in the second centimetre of the wood by November were similar, but smaller (Fig. S3). Furthermore, the observed treatment effects on wood nonstructural carbon concentrations were comparatively small in relation to the seasonal variation (Fig. 6). With the exception of a decrease in root starch of 0.65 ± 0.17% (t = −3.83) relative to control, nonstructural carbon concentrations were similar in tissues below the treatments.

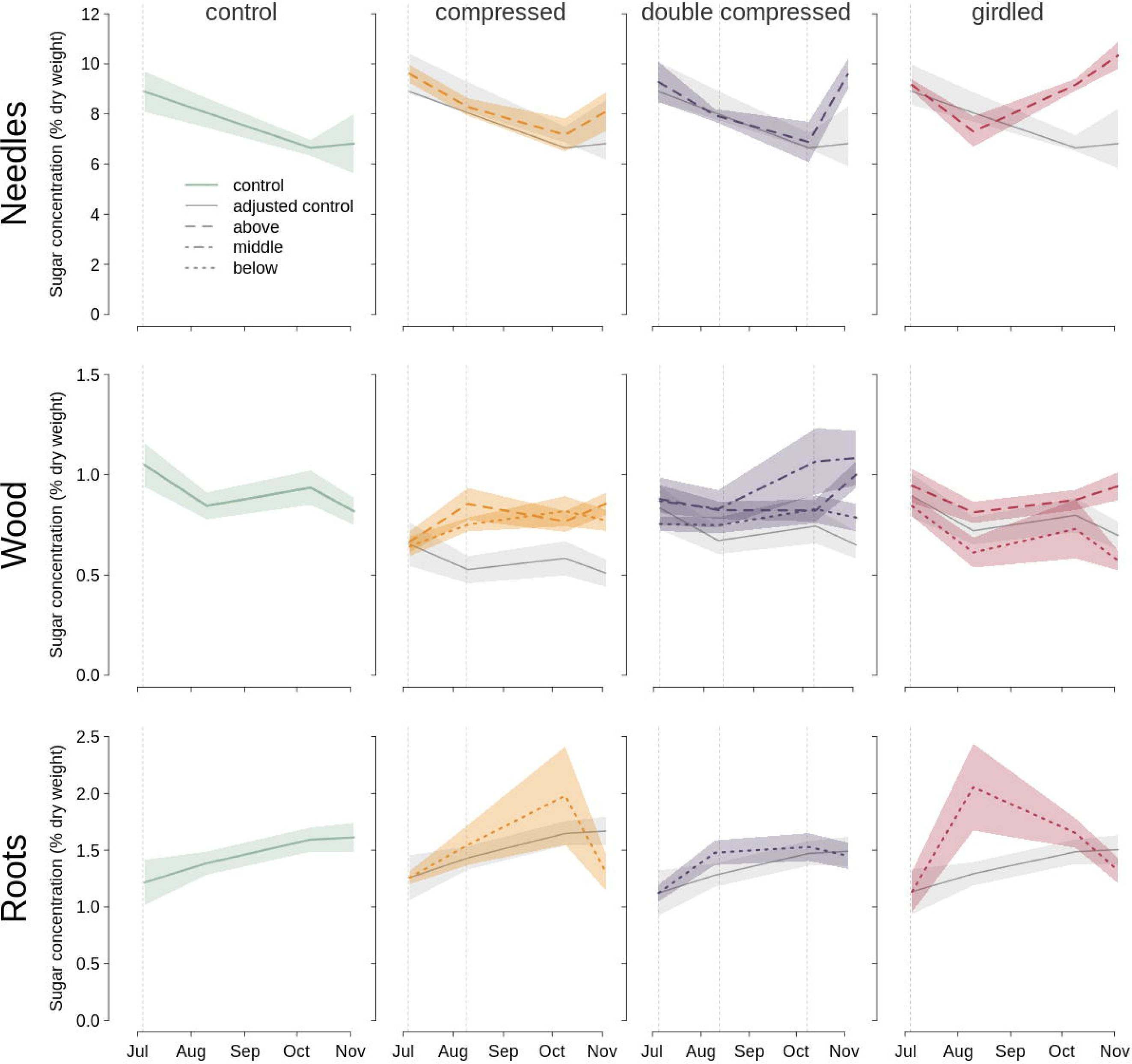

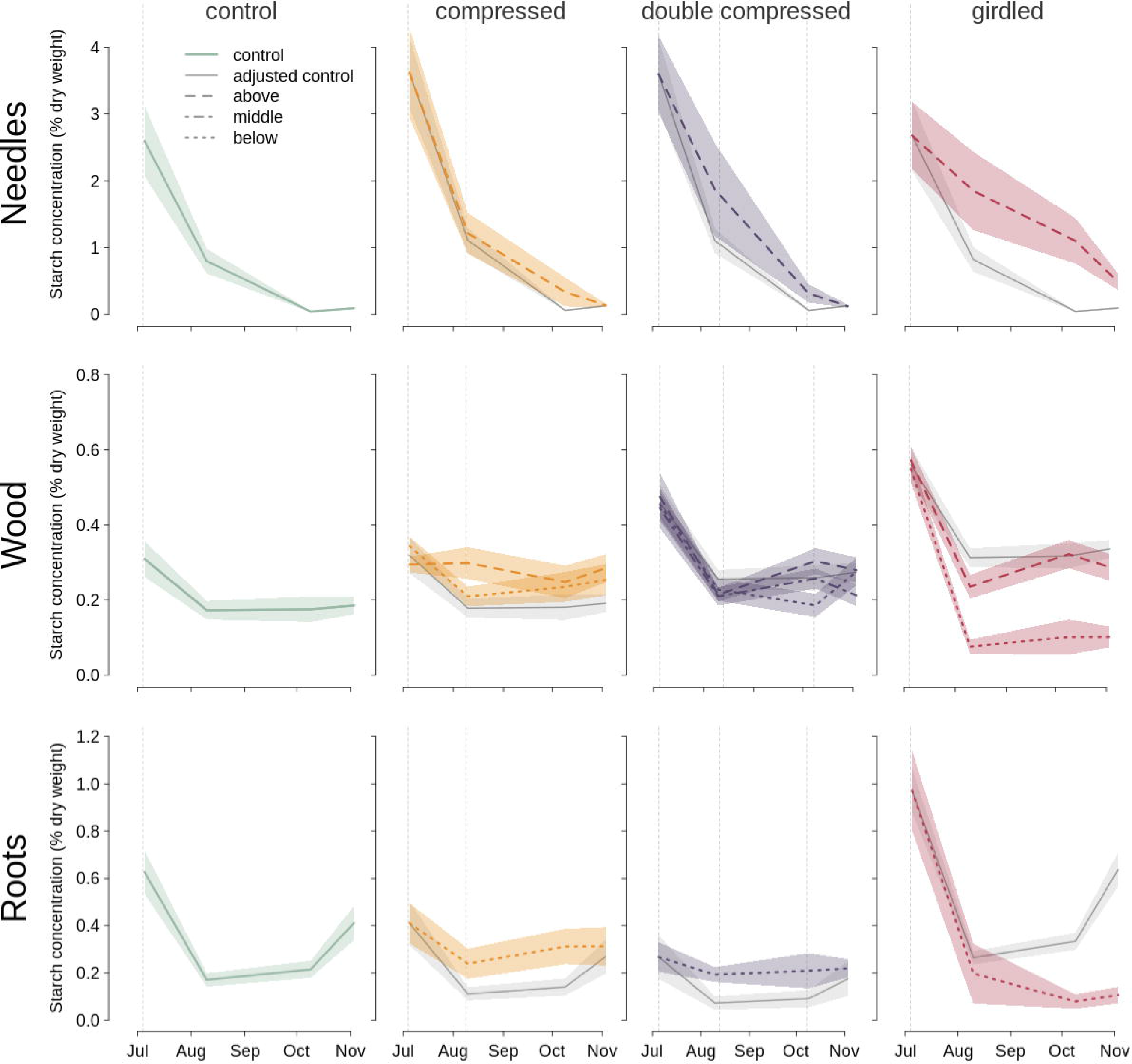

## 4. Discussion

Our observations suggest that phloem compression was successful as an alternative to girdling to reduce phloem flow. Similar to Henriksson *et al*. (2015), we observed diverging CO_2_ efflux rates above and below compression treatments. However, convergence of rates above and below the compression after removal of compression treatments took several months here, while it happened within weeks for Henriksson *et al*. (2015). The growth increase, especially above the double compression treatment, further suggests that the treatment generated an effective bottleneck for phloem transport, leading to enhanced carbon and/or hormone supply above the treatment. However, the small increase above the single compression and similar growth below the compression treatments relative to the control, without a substantial depletion of connected nonstructural carbon reserves, suggests that phloem flow was only reduced and not halted completely. Wood also continued to form between the double compression collars with CO_2_ efflux rates roughly equaling control group rates. The maintenance of growth between the compression collars happened without reducing the local stem nonstructural carbon reserves (first and second centimetre), indicating that some carbon continued to be transported across the compression zones. Furthermore, each compression zone enhanced the difference in CO_2_ efflux in the double compression, which could also be a result of leakage across each compression zone. Because phloem compression was effective, albeit somewhat leaky, phloem transport seems to have been successfully modified to generate our hypothesised carbon-supply gradient ranging from severe carbon limitation below the girdle, over moderate carbon limitation below the compression (due to some leakage) and moderate carbon supply surplus above the compression, to a larger increase of carbon supply above the girdle. Phloem transport is a mass flow. Consequently, simultaneous reduction in other phloem-transported signaling compounds, which are essential for wood formation (Aloni, 2013; Buttò *et al*., 2020), could occur and may have influenced our observed results.

Multiple studies using girdling have shown that stopping phloem transport causes an early cessation of cambial activity (Maunoury-Danger *et al*., 2010; Oberhuber *et al*., 2017). Our results showed that new cells formed below both compression treatments after removal in the same (20 out 20 trees) and the following (19 out 20 trees) growing season, indicating that the phloem can recover from compression and carbon and/or hormone availability can re-activate the cambial meristem once the phloem has recovered. A similar number of cells formed below the single compression relative to control after the removal of the compression, but slightly less cells formed below the double compression (possibly due to later release). Mass growth after the removal of the compression was on par with the control group, suggesting that growth resumes at normal levels and there is no compensatory enhancement of growth. While our sampling frequency did not allow for the precise identification of critical dates of wood formation, our results show that cambial activity is dependent on intact phloem transport and that compressed phloem tissue can partially recover as late as close to the end of the growing season (here early October). Nevertheless, the treatment had lagged effects on wood formation in compressed trees in the following season, suggesting that full recovery takes more than one growing season. Phloem compression appears to be an exciting new tool to help to understand which physiological processes may be carbon-supply limited, over which timescales, and during what time of the year.

Growth and CO_2_ efflux covaried with the presumed carbon-supply rate, but the size of nonstructural carbon pools did not change substantially, illustrating the allocation of carbon to growth, CO_2_ efflux, and nonstructural carbon reserves is dynamic under varying carbon supply. Under low carbon supply, a larger fraction of carbon was allocated to maintain nonstructural carbon pools close to control group values, resulting in only marginal net declines in nonstructural carbon pools. A similar prioritisation of nonstructural carbon reserves was observed under low carbon supply due to defoliation (Piper *et al*., 2015; Wiley *et al*., 2017), drought (Hagedorn *et al*., 2016), and low atmospheric CO_2_ (Hartmann *et al*., 2015). The fraction of carbon invested in respiration and woody tissues was higher under elevated carbon supply compared to the control. This contrasts with whole-tree effects of proportionally larger increases in nonstructural carbon concentrations than stimulations of wood growth under elevated CO_2_ (Ainsworth and Long, 2005; Körner *et al*., 2005). Other direct phloem manipulations support our observed shift in allocation towards growth at elevated carbon supply above the treatment (Maier *et al*., 2010; Regier *et al*., 2010; De Schepper *et al*., 2011; Oberhuber *et al*., 2017). The apparent discrepancy may result from within tree feedbacks at elevated CO_2_, such as non-stomatal downregulation of photosynthesis (Salmon *et al*., 2020), re-distribution of carbon throughout the entire tree, or difference in signalling. Importantly, we found that carbon consumption by wood growth and CO_2_ efflux are much more sensitive to variations in cambial carbon supply than are nonstructural carbon pools.

### 4.1 Wood growth was correlated with carbon supply

Our first hypothesis (H1) was only partially supported, because only the number of newly formed cells but not mean cell-wall area or mean cell size was strongly correlated with carbon supply. While detecting strong trends of decreased mean cell-wall area at low carbon supply was complicated by the reduced number of cells formed, cell size was unaffected by carbon supply. The number of cells that formed after treatment onset was strongly related to the presumed carbon supply. Availability of soluble sugars is linked to cell division in plants through metabolica signalling (Smith and Stitt, 2007; Lastdrager *et al*., 2014). Soluble sugar concentrations also influence the osmotic potential in cambial cell lumens which affects turgor pressure (Guerriero *et al*., 2014), hence growth (Peters *et al*., 2020). Previously, modelling (De Schepper and Steppe, 2011) and experimental evidence from saplings (Winkler and Oberhuber, 2017) was interpreted to suggest that an accumulation of osmotically active sugars in the cambium was responsible for observed increases in cell division due to consequent changes in turgor pressure. However, we do not see an increase in soluble sugar concentrations in the first or second centimetre of the xylem, despite large increases in the number of cells formed. Due to strong radial gradients in soluble sugar concentrations in the cambial zone (Uggla *et al*., 2001), and our comparatively coarse measurement resolution (e.g., 1cm) we cannot rule out turgor-mediated mechanisms causing the observed correlations. Alternatively, soluble sugars could be a signal directly operating on cell division (Riou-Khamlichi *et al*., 2000). Dynamic soluble sugar concentrations have already been argued to drive the early-to-latewood transition (Cartenì *et al*., 2018), which is supported by the observed decline in mean cell-wall area at low carbon supply here. While limitation in carbon-supply due to natural defoliation can reduce growth (Fierravanti *et al*., 2019) and cell numbers substantially (Castagneri *et al*., 2020), we found that the cumulative number of cells formed, thus cell division, and to a smaller degree cell-wall deposition, seem to be regulated by carbon supply more generally (including at elevated carbon supply). Interestingly, cell size was not affected, despite the marked increase in cell number above all phloem transport manipulations, resulting in much higher cumulative cell-wall area, thus biomass.

Contrary to previous studies, we did not see a reduction in cell-wall area above any of the treatments. Reduced cell-wall thickness in latewood has been reported for Norway spruce saplings above the girdle (Winkler and Oberhuber, 2017). This reduction in cell-wall deposition at higher carbon supply was attributed to either additional carbon demand due to more cells formed or alternative investment in defense compounds due to a wound reaction. Winkler & Oberhuber (2017) also reported the formation of smaller lumina in earlywood above the girdle and larger lumina in latewood, in contrast to our findings of no substantial changes in cell size and wall area. We cannot rule out that differences in phenology, especially because earlywood formation was complete at the onset of our experiment, or species may be responsible for the differing effects on cell-wall deposition and cell size. We also acknowledge that Winkler & Oberhuber (2017) measured cell-wall thickness and lumen area, while we present data on cell-wall area and cell diameter. Nonetheless, we suspect that the differences are caused by different developmental stages of the trees: saplings versus mature trees, as trees are thought to be more carbon limited at younger ages (Körner, 2003; Hayat *et al*., 2017; Hartmann *et al*., 2018). Importantly, both results indicate that higher carbon seems to trigger the proliferation of additional cells and investment of carbon into cell walls of additionally formed cells, instead of thickening cell-walls of normally formed cells.

### 4.3 CO_2_ efflux covaried with carbon supply

Stem CO_2_ efflux rates increased with presumed carbon supply, supporting our second hyposthesis (H2) that the two will covary. CO_2_ efflux rates responded within weeks to phloem transport manipulations, but took substantially longer to relax to control group values once the compression was removed. The only other stem-compression study documented a similar effect on stem CO_2_ efflux, but a faster recovery after only two weeks for mature Scots pines (Henriksson *et al*., 2015). We suspect that this discrepancy in recovery is caused by our experiment starting later in the growing season, thus allowing only a smaller opportunity for re-growth and/or recovery of the phloem. The compression collar design used here also exerted slightly higher pressure around the circumference (Henriksson and Rademacher, 2019), which may have contributed to the longer observed recovery times and continued effects in double compressed trees on radial growth in the following growing season.

Stem CO_2_ efflux declined as much below compression treatments as below girdles, despite mass growth remaining at control group levels below both compression treatments. CO_2_ efflux may simply be more sensitive to reduced carbon supply. The continuation of growth below the compression treatments seems to suggest that non-growth metabolism is preferentially downregulated at lower carbon supply. CO_2_ efflux and growth may also preferentially draw on different carbon sources (i.e., phloem-transported versus local stores). Indeed, isotopic studies have revealed that respiration preferentially uses younger carbon, while growth draws preferentially on older carbon from reserves (Maunoury-Danger *et al*., 2010). Similar CO_2_ efflux levels below the girdle and both compression treatments suggest that they both approached a necessary minimum that is essential to maintain living tissue (Minchin and Lacointe, 2005). Because wood formation commits more than the instantaneously required resources, it is reasonable to assume that respiration is preferentially down-regulated to maintain reserves needed to fuel and provide resources for cell expansion and cell-wall thickening of cells that have just divided. Given the lack of local depletion of nonstructural carbon reserves below treatments, we assume the majority of the carbon necessary to fuel CO_2_ efflux was supplied from root reserves in the girdled trees and from leakage across the compression zone in compressed trees.

### 4.4 Wood soluble sugar and starch concentrations were stable across a large carbon-supply gradient

Contrary to our third hypothesis (H3), soluble sugar concentrations remained remarkably constant for any individual tissue among treatments. One exception was increases in needle soluble sugar concentrations in girdled trees, which may cause non-stomatal photosynthetic down-regulation (Salmon *et al*., 2020) and more generally trigger whole-plant feedbacks. In contrast to stable sugar concentrations, starch concentrations varied in a few tissues, potentially to buffer soluble sugar concentrations against imbalances in carbon supply and demand. Noticeable remobilisation and accumulation of starch was apparent in the roots and needles of girdled trees, respectively, and to a lesser degree in the double compression. Overall, the net changes of nonstructural carbon reserves were marginal compared to investments in growth and losses to CO_2_ efflux.

The observed stable soluble sugar concentrations across treatments add to a growing body of evidence that nonstructural carbon concentrations follow relatively constrained seasonal cycles in the wood of mature trees (e.g., Zhang *et al*., 2020). This maintenance of a seasonal rhythm in soluble sugar concentrations in the cambium suggests that equating high wood soluble sugar concentrations with a sink limitation (Hagedorn *et al*., 2016) is flawed, because growth and wood soluble sugar concentration seem to be regulated independently under varying carbon supply.

### 4.5 Conclusion

Our study has demonstrated that stem compression is an effective and reversible way to manipulate phloem flow, which can be harnessed to study the temporal effects of variation in phloem transport and its recovery. Restricting phloem transport by compressing and girdling trees has illustrated homeostatic maintenance of soluble sugar concentrations along a presumed large gradient of carbon supply, whereas wood growth and CO_2_ efflux covaried with carbon supply. Concerning wood formation, cell division seems particularly sensitive to carbon supply with minor additional effects on cell-wall deposition. Overall, wood formation seems to be directly carbon-limited.

## Acknowledgements

AF, AR, TR, and YC acknowledge support from the Natural Environment Research Council (NE/P011462/1) and National Science Foundation (DEB-1741585). AR is also supported by the National Science Foundation under grant DEB-1237491 and DEB-1832210. DB acknowledges support through the Swiss National Science Foundation (PSBSP3-168701) and the Harvard Forest Bullard Fellowship. We also thank Aglaé Landry-Boisvert, Brooklynn Abaroa, Emory Ellis, Kyle Wyche, and Mark VanScoy for help in the field, Shawna Greyeyes, Amberlee Pavey, and Angelina Valenzuela for help in the lab, Katharyn Duffy, Teemu Hölttä, Drew M. P. Peltier, and three anonymous reviewers for feedback on the manuscript, and Henrik Hartmann, Nils Henriksson, Teemu Hölttä, and Cyrille Rathgeber for friendly peer-review of the ideas and methods.

## Author Contributions

TR and AR planned and designed the experiment. TR conducted the experiment, collected all data and materials. TR, JL, MF and PF conducted the laboratory analyses. BS, DB and TR created the TRIAD platform. TR developed the NSCprocessR package with input from JL and AR. TR performed the statistical analysis, generated the figures, and wrote the paper. All co-authors discussed ideas, provided feedback, edited the manuscript draft, and approved the manuscript for submission.

