## Supplementary Information 1 for "Manipulating phloem transport affects wood formation but not nonstructural carbon concentrations in an evergreen conifer"

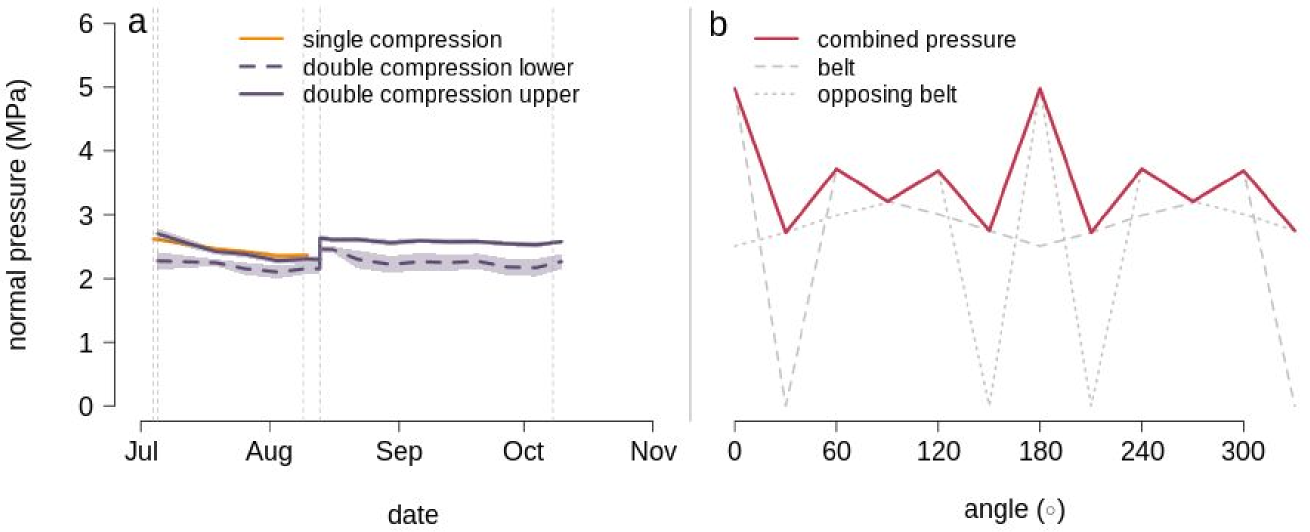


Fig. S1 - (a) Mean collar pressure and one standard deviation (shading) over time for the single (orange) and double compression (purple) treatments. Key dates (e.g., start, re-tightening and end) are indicated by the grey dashed vertical lines. (b) Circumferential pressure distribution as measured in the lab before deployment using piezo-electric pressure sensors. The dashed and dotted lines show the pressure exerted by an individual compression belt, and the red line the combined pressure of two diametrically opposing belts.
