## Supplementary Information 2 for "Manipulating phloem transport affects wood formation but not nonstructural carbon concentrations in an evergreen conifer"

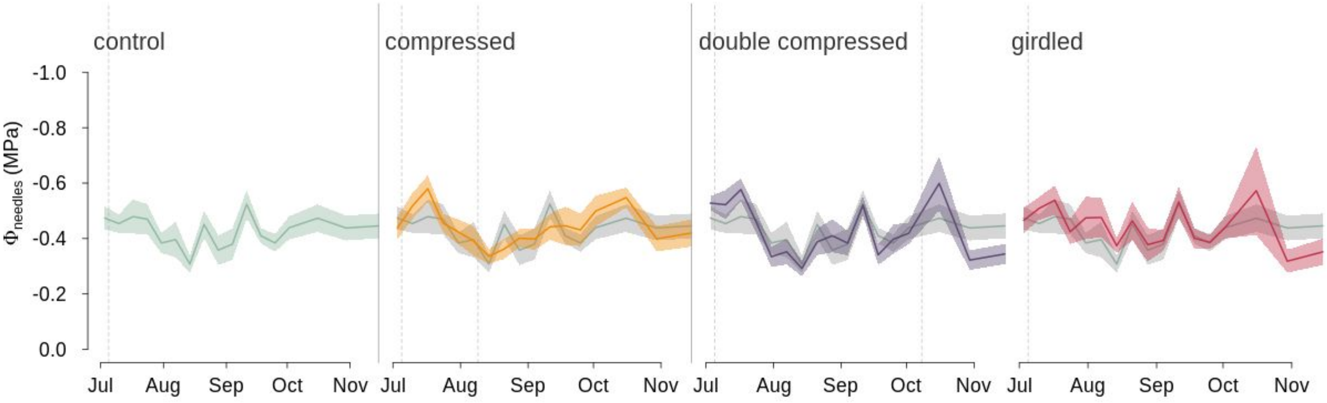


Fig. S2 - Weekly mean (solid line) and one standard error (shading) of needle water potential ($\Phi_{needles}$) for the control, compressed, double-compressed, and girdled trees. The control group mean and standard error are shown on treatment panels in grey to facilitate comparison. Key dates for each treatment (e.g., start, re-tightening and end date) are indicated by dashed grey vertical lines. Pre-dawn needle and branch water potential were measured with a pressure chamber (Model 600, PMS Instruments, Albany, Oregon, USA) once per week per tree from the end of June to the beginning of November (displayed above) and twice in the two following years (not shown).

For each measurement, two branch tips per tree were cut, sealed in air-tight plastic bags for transport. Initial tests showed that water potential could be determined accurately up to 20 minutes after cutting the tissue. Within 20 minutes of harvest, water potential was determined after re-cutting the tissue. Observed needle and branch water potential did not differ among treatments during the duration of the experiment or in the remaining growing season.

After about one year, the girdled trees had substantially more negative needle water potentials (not shown), from -046 ± 0.06 MPa to -0.91 ± 0.08 MPa (5.20). All girdled trees showed clear signs of decline, such as canopy browning, and seven trees died about two years after the girdling (2019). With the exception of one compressed tree, which died in 2018, neither the single nor the double compression had a detectable effect on pre-dawn water potentials for the two following growing seasons.
