## Supplementary Information 3 for "Manipulating phloem transport affects wood formation but not nonstructural carbon concentrations in an evergreen conifer"

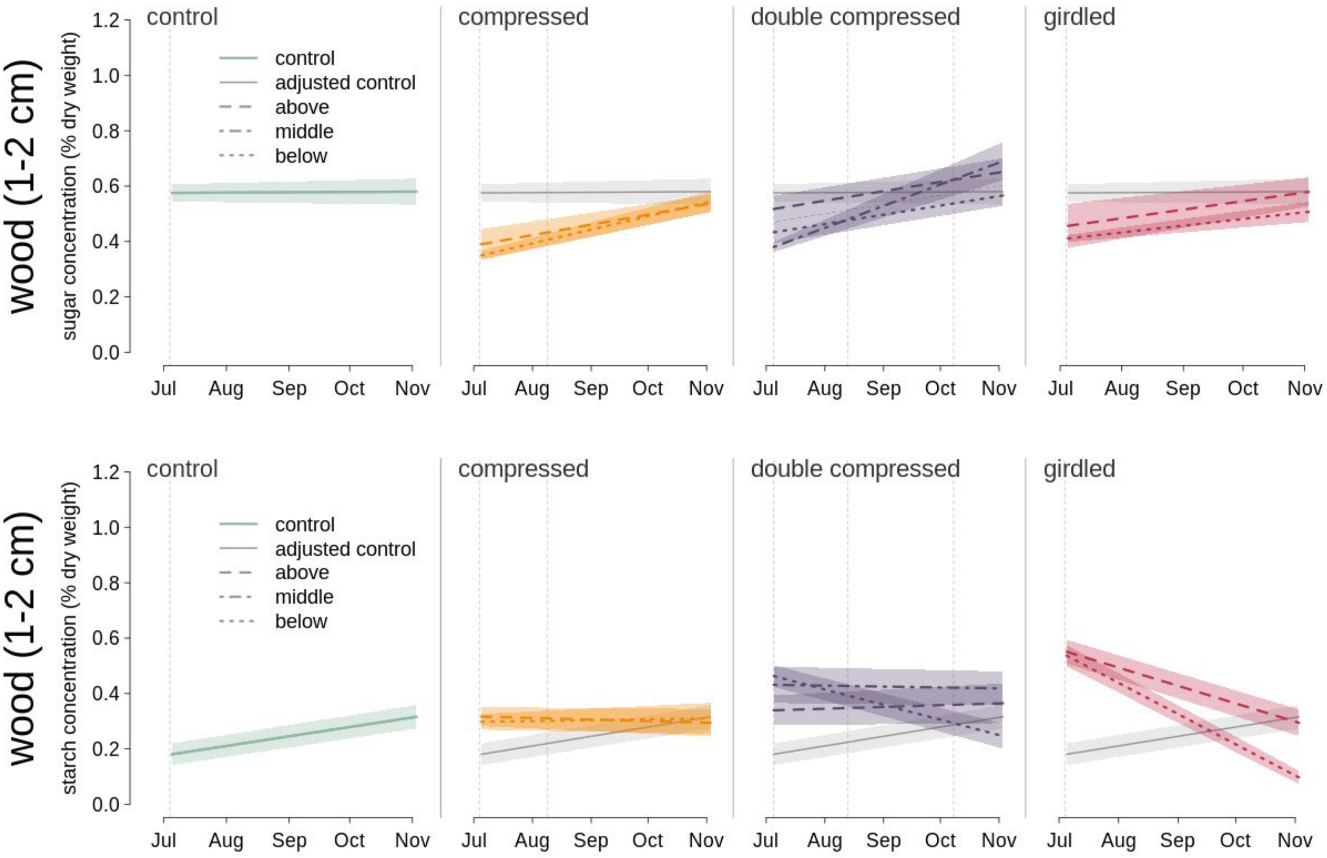


Fig. S3 - Soluble sugar (top row) and starch concentrations (bottom row) in the second centimetre (e.g., 1 to 2 cm range) of the wood for the control (left column; green), compressed (second column from the left; orange), double compressed (second column from the right; purple), and girdled trees (right column; red). The lines indicate the mean concentration with shading representing one standard error. For ease of comparison, control values were added to all treatment panels in grey. Key dates for each treatment (e.g., start, re-tightening and end date) are indicated by dashed grey vertical lines. Dashed lines correspond to tissues above the treatment, dotted line to tissue below the treatment and dashed-dotted lines to tissues in the middle of the double compression.

The changes in concentrations from July to November were less than in the first centimetre (t-test, $p = 0.0073$ for sugar and $p = 0.0004$ for starch).
