## Supplementary Information 4 for "Manipulating phloem transport affects wood formation but not nonstructural carbon concentrations in an evergreen conifer"

### Supplementary Information 4: Scaling structural carbon

Tim Rademacher

13/11/2020

*This pdf was generated from an Rmarkdown file, which includes all R code necessary to reproduce the estimatations. The Rmarkdown file is available on github: https://github.com/TTRademacher/Exp2017Analysis.*

#### Structural carbon estimation for stem sections

We estimate structural carbon gain for each stem section and period from the cell-wall area estimated to have formed in that period. The cell-wall area of one thinsection sampled at the end of the growing season ($3^{rd}$ November 2017) was therefore divided into sections grown between sampling dates (e.g., prior, during and after treatment) by association with the fraction of the ring formed at each sampling date. For each sampling date two microcores were collected with trephor tool (Rossi et al., 2006). Microcores in August, October and November were punched in close proximity to the July sample. However, the samples were offset by, at least, 5cm horizontally on the same vertical axis to avoid sampling a wound reaction of the injury inflicted by previous cores. As growth varies locally around the circumference of the stem, sampling relatively close was a first step to minimised local differences in growth between sampling dates. One of those microcores was embedded in parafin (Tissue Processor 1020 Leica, Switzerland) and 7 $\mu m$-thick crossectional cuts were obtained using a rotary microtome (Leica RM2245, Switzerland). The cuts were stained with astra blue and safranin red, and scanned on a sliding scanner (resolution of roughly 1.5 $\mu m$). Where the resulting image of the first microcore did not fully include the necessary rings or had other quality issues, we cut and processed the second microcore. Between the two microcores, we were able to obtain high-quality image data for 315 out of 320 samples (i.e., 4 sampling dates times 40 trees times on average two sampling heights). Only for one tree (tree id: 06; treament: control) both thinsections collected in October did not contain a full set of rings necessary for the analysis. For each image, we identified and divided the ring formed in 2017 into 20 $\mu m$-wide discrete tangential bands and derived the average anatomical structure for cells in each band using ROXAS (Arx and Carrer, 2014). In particular, we measured average cell-wall area, cell-wall thickness, as well as tangential and radial lumen diameter. The ring width of the fully formed 2017 ring ranged between 0.42 and 3.23 mm with a mean of 1.64 mm. Even the smallest ring was divided into 21 discrete bands of a width of 20-$\mu m$ with associated anatomical data.

##### Estimating period of formation of tangential bands

Ring widths were measured for all sampling dates using the self-developed Wood Image Analysis and Database plaform (http://wiad.science). The ring widths were standardised to account for local variations in growth in thinsections. Standardised ring widths from the different sampling dates were then used to associate proportions of the final ring from the November microcore (i.e., which $20 - \mu m$-wide band to include), to specific growth periods. For this purpose we identified and measured ring widths in thinsections taken prior (e.g., $3^{rd}$ of July 2017) and during the treatment (e.g., $10^{th}$ of August 2017, and $9^{th}$ of October 2017), as well as at the end of the growing season ($3^{rd}$ of November). For standardisation, we used the width of an older ring from the same thinsection to correct for local variation in growth between microcores.

$$f_{m,17} = \frac{rw_{m,17} \times rw_{end,15}}{rw_{m,15}} \times \frac{1}{rw_{end,17}}$$

, where $f_{m,17}$ is the fraction of the ring width formed in month $m$ (e.g., July, August, or October), $rw_{m,15}$ and $rw_{m,17}$ are respecitvely the 2015 and 2017 ring width in the thinsection collected in month $m$, and $rw_{end,15}$ and $rw_{end,17}$ are respectively the 2015 and 2017 ring widths in the thinsection from the November sample. This assumes that the ring was fully formed on the $3^{rd}$ of November, which is about two weeks after the typical end of cell wall thickening for white pine at Harvard Forest. Then, $f_{m,17}$ theoretically varies between 0 and 1 with 0 meaning none of the ring was formed at the sampling date and 1 meaning that the entire ring was already formed.

The number of measurements for growth rings drops off strongly after the 2015 growth ring (table 1), because the older rings are further away from the bark, hence they are less likely within the 15mm reach of the trephor tool (Rossi et al., 2006) used to collect the microcores. Among the recent years with high replication, 2015 was preferred over 2016, because of an abnormally dry growing season in 2016 (total annual precipitation of 936 $mm$, which is 160 below the long-term average of 1096 $mm$). In fact, about three quarters of trees included in this study showed an intra-annual density fluctuation in 2016 due to the dry growing season and the remaining quarter of trees had a relatively narrower ring. 2016 was 0.90 °C warmer with 234 mm less precipitation compared to the 30-year average. Consequently, standardisation by the 2016 ring would introduce a systematic bias based on the bi-modal ring width distribution in 2016. In contrast, 2015 did not include any outstading climatological events or resulting systematic bias in growth. Indeed, the 2015 growing season climate was similar to the 2017 growing season climate with a mean annual temperature of 8.1 $°C$ and a total annual precipitation of 1024 $mm$. Both 2017 and 2015 were close to the long-term averages mentioned above.

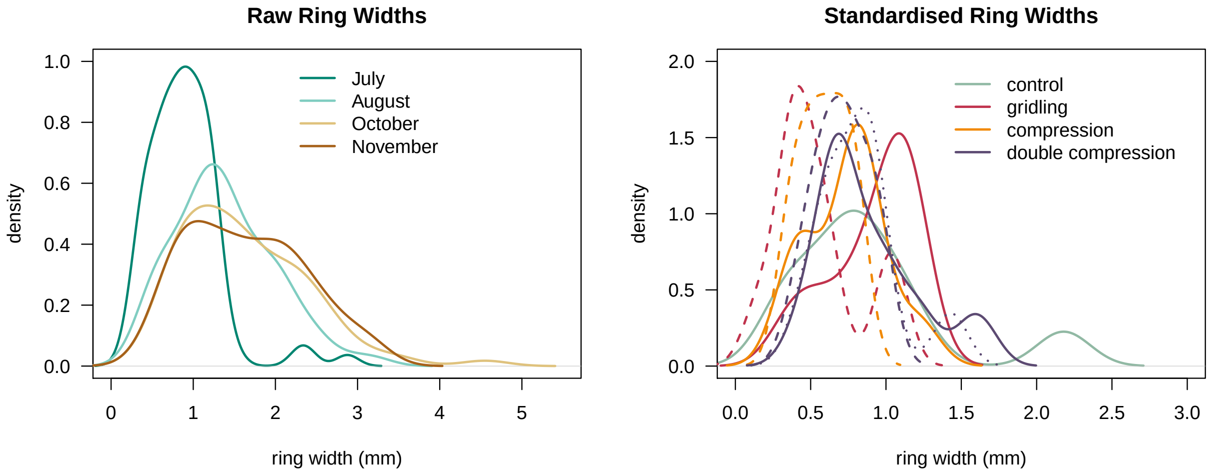


Table 1: Number of measured growth ring widths for all trees out of a potential 320 thinsection images.

| | 2017 | 2016 | 2015 | 2014 | 2013 | 2012 | 2011 | 2010 | 2009 | 2008 |
|---|---|---|---|---|---|---|---|---|---|---|
| Number of Samples | 320.00 | 319.00 | 315.00 | 282.00 | 209.00 | 115.00 | 49.0 | 31.0 | 9.0 | 0 |
| Percentage | 100.00 | 99.70 | 98.40 | 88.10 | 65.30 | 35.90 | 15.3 | 9.7 | 2.8 | 0 |
| Mean Ring Width (mm) | 1.71 | 1.83 | 2.35 | 2.07 | 1.55 | 1.85 | NaN | NaN | NaN | NaN |

On average, 55%, 83.7%, and 101.6% of the ring width had respectively grown in July, August and October. Thus, a bit more than half of the ring widths was already formed before the start of the treatment and ring width growth had *de facto* finished by Ocotober. However, there were substantial differences between treatments and sampling heights in the timeline of xylogenesis.

Using the fractional ring widths as boundaries, each 20-$\mu m$ band was associated with one of the four periods: (1) beginning of growing season to start of experiment ($3^{rd}$ of July 2017), (2) first month of the experiment ($4^{th}$ of July to $9^{th}$ of August 2017), (3) second month of the experiment ($10^{th}$ of August to $8^{th}$ of October 2017), (4) end of double compression to end of growing season ($9^{th}$ of October to $3^{rd}$ of November 2017). These period are illustrated in Fig. 1 (main manuscript).

##### Scaling measured cell wall area to stem section

After we associated each tangential band with a growth period, each band is scaled to the circumference of the stem section by multiplication of the mean zonal cell-wall area by the number of times the average zonal tangential cell width ($w_{t,s}$) fits into the entire circumference of each stem section. To account for the bark (assuming a bark thickness of 0.8cm) we reduced the circumference, as measured at the beginning of the experiment ($CBH_{outer,s}$) according to the following equation:

$$CBH_s = CBH_{outer,s} - 0.8cm \times 2\pi$$

The zonal cell wall area for each band is then summed across fractions of the ring were previously associated with the same period of formation. As a result we obtain an estimate of the total cell wall area formed during each period.

To further scale the total circumferential cell-wall area to a cell-wall volume per stem section, we assume that the measured cell-wall area ($CWA_{s,p}$) is representative of the longitudinal wood profile for the stem section, hence we can simply multiply it with the height of the stem section (h = 10cm).

$$V_{CW,s,p} = \frac{CBH_s}{w_{t,s}} \times h_s \times C\bar{W}A$$

Implicitly, this approach assumes that wood is circumferentially and longitudinally homogenous and the measured tracheids are representative of the woody tissue. thus, the approach does not account for other types of tissue than tracheids, such as rays and resin ducts. Tracheids make up 93% of the volume of wood in white pines with resin ducts and rays accounting for 1% and 6%, respectively (Brown et al., 1949). However, since there was no clear treatment effect on the occurance of resin ducts or rays cells, this assumption should not introduce a bias in carbon sequestration in strucutre between treatments. As ray cells and resin ducts have a slightly lower carbon density, this approach does, however, over estimate carbon density marginally.

Finally, the cell-wall volume for each ring portion of the stem section ($V_{CW,s,p}$) is multiplied by a fixed cell-wall density ($\rho_{CW}$) of 1.489 $kg\,m^{-3}$ (measured for Scots pine by Plötze and Niemz (2011)) and a carbon content of 49.74% typical of white pine (Lamlom and Savidge, 2003) to estimate the total structural carbon gain per stem section and period ($M_{p,s}$ in g).

$$M_{s,p} = V_{CW,s,p} \times \rho_{CW} \times 49.74\%$$

##### Alternative estimate using wood density

To better understand uncertanties associated witgh our assumptions we also used an alternative estimate based on wood density. For this alternative estimation, the carbon is simply estimated as a product of the volume of wood and an average wood density. To obtain wood density values, X-ray densiometry was performed on 18 increment cores from the ten control trees.
