## Supplementary Information 5 for "Manipulating phloem transport affects wood formation but not nonstructural carbon concentrations in an evergreen conifer"

### Supplementary Information 5: Estimating losses due to $CO_2$ efflux

Tim Rademacher

13/11/2020

*This pdf was generated from an Rmarkdown file, which includes all R code necessary to reproduce the estimatations. The Rmarkdown file is available on github: https://github.com/TTRademacher/Exp2017Analysis.*

#### Estimating total carbon losses due to $CO_2$ efflux

To estimate stem carbon losses due to $CO_2$ efflux, $CO_2$ efflux measurements were made weekly using a Li-Cor820 starting at 13.00h. The LiCor820 was attached to a PVC pipe (4" diameter) that were previously fitted and attached to the stem sections using silicone to create a closed chamber in which the enclosed air circulates via a small pump, similar to the Flux Puppy system (Carbone et al., 2019). $CO_2$ efflux and uncertainties were calculate from the change in $CO_2$ concentrations using the RespChamberProc package (Perez-Priego et al., 2015) in R (R Core Team, 2019). Stem sections were visited in the same order to reduce the role of diel variations over the course of the experiment. The order was initially randomised but slightly adjusted to assure that no treatment was systematically measured later in the day. Stem $CO_2$ efflux is also related to temperature and water availability (Yang et al., 2016). Air temperatures and volumetric soil water content near the site ranged between 7.1 and 23.9°C and 11.3 and 44.8% over the experimment, but less than 2.9°C or 0.1% on any particular measurement dperiod (Boose and Gould, 2019). All data is publicly available on the Harvard Forest Data Archive (Rademacher and Richardson, 2020).

#### Integration over the surface area of each stem section

Instantaneous fluxes of diffusive $CO_2$ loss through the bark were measured as described above and subsequently integrated across the surface area of each stem section to estimate the respiratrory loss in grams per stem section per day. To integrate the measured $CO_2$ efflux rates across the surface of each 10cm-stem section, we first calculated the surface area of the i-th stem section ($A_{s,i}$) as follows:

$$A_{s,i} = \frac{cbh_{s,i}}{100} \times h$$

, where is the $cbh_{s,i}$ is the circumference in cemtimetres, as measured in the field with a tape measure, and $h$ is the height of the section in metres (here $h = 0.1$m). The stem $CO_2$ efflux rate over this surface area ($R_{i,k}$) was then determined by multiplication of the instanteanous $CO_2$ flux ($f_{i,CO2}$; in $g\,m^{-2}\,day^{-1}$) and the surface area to get the $CO_2$ flux rate (in $g\,day_{-1}$).

$$R_i = f_{i,CO2} \times A_{s,i}$$

#### Estimate respiratory losses for each periods

$CO_2$ flux rates were averaged for over four time periods sensu Figure 1 of the main text: (0) before experimental onset, (1) first month after start of the experiment, (2) second and third month after the start of the experiment, and (3) from the fourth month after the start of the experiment to late autumn. We then approximated the total loss of carbon for each combination of period $j$ and stem section $i$ ($R_{i,j}$) by multiplying the temporal mean of spatially integrated $CO_2$ flux rates ($R_i$; in $g\,day^{-1}$) by the length of the period ($l_p$ in days).

$$R_{i,j} = \frac{1}{n}\sum_{k=1}^{n} R_{i,k} \times l_p$$

, where n is the number of weekly sampling dates during each period. The figure below shows the average and standard deviation of the mean loss of carbon due to $CO_2$ efflux for each period (i.e., above the start date of each period).

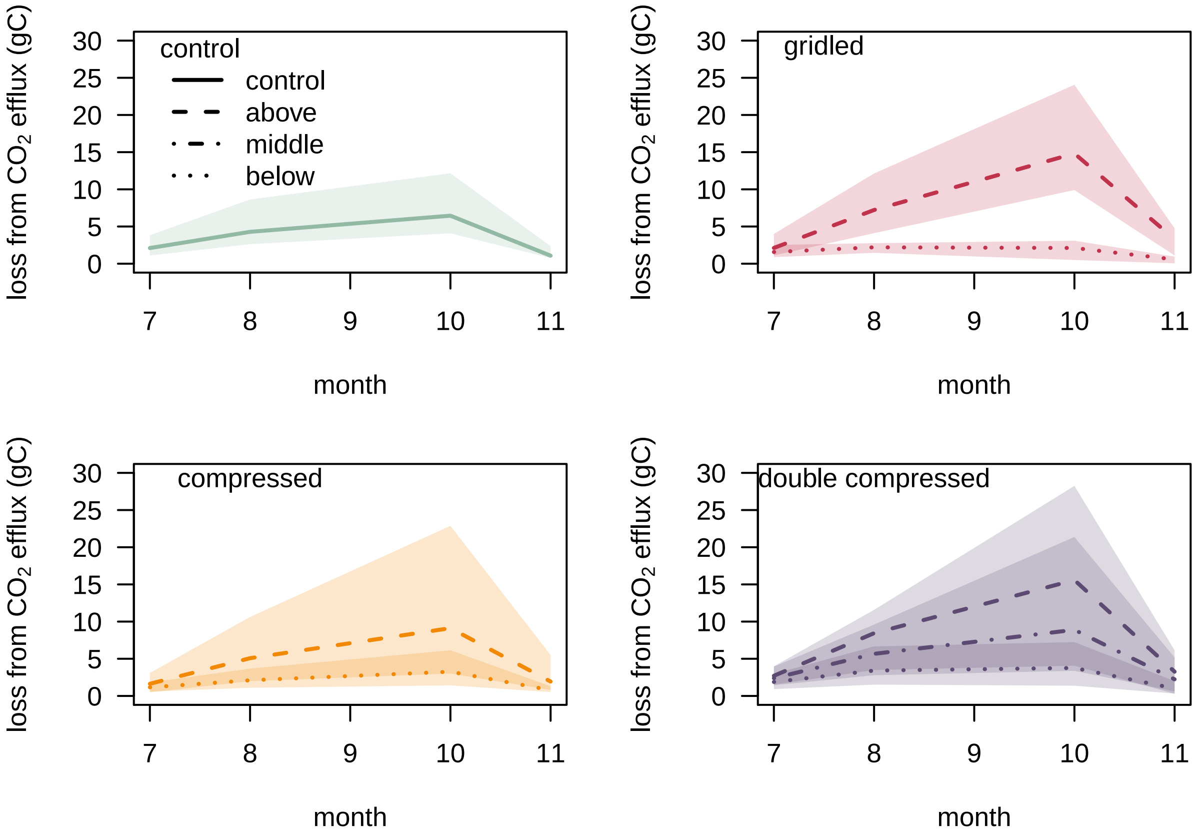
