## Supplementary Information 6 for "Manipulating phloem transport affects wood formation but not nonstructural carbon concentrations in an evergreen conifer"

### Supplementary Information 6: Scaling net changes in nonstructural carbon reserves

Tim Rademacher

13/11/2020

*This pdf was generated from an Rmarkdown file, which includes all R code necessary to reproduce the estimatations. The Rmarkdown file is available on github: https://github.com/TTRademacher/Exp2017Analysis.*

#### Estimating total nonstructural carbon reserves and changes over time

To estimate changes in soluble sugar and starch (measured as glucose equivalent) for each stem section, we scaled the concentrations derived using colorimetric assays, as described in the main text, to the total mass of easily accessible nonstructural carbon reserves including sugar and starch per stem section. Because reserves in deeper tissues are generally smaller (Furze et al., 2019; Hoch et al., 2003), are assumed to be less accessible (Carbone et al., 2013), and each sample's analysis requires substantial effort in the lab, we limit our analysis to the first two centimetres. The first two centimetres underlying the bark for each stem section will constitute the largest and most easily accessible amount of the radial nonstructural carbon reserves, as nonstructural carbon concentrations follow a generally decreasing gradient from the bark towards the pith (Furze et al., 2019; Hoch et al., 2003). Indeed, the second centimetre already showed similar, yet substantially smaller changes from July to November (Fig. S3) than the first centimetre. Below, we describe the exact methods for estimating the easily accessible total nonstructural carbon reserves in stem sections (i.e., 0-2cm of radial profile). Net differences in these total reserves over time (i.e., four sampling dates sensu Fig. 1 of main text) provide estimates of net gains or losses of easily accessible total soluble sugar and starch mass for each stem section (Fig. 2 of main text). Below we provide exact methods and an example calculation. All data is publicly available on the Harvard Forest Data Archive (Rademacher and Richardson, 2020).

Initially, we measured nonstructural carbon concentrations for the first and second centimetre of each stem core as detailed in the main text. We then derive the total mass of nonstructural carbon in sugars and strach ($M_{sugar,i}$ and $M_{starch,i}$, respectively) by multiplying the dry matter of the hollow cylinder ($M_i$) of each stem section $i$ with the respective section's sugar and starch concentrations ($c_{sugar,i}$ and $c_{starch,i}$). For sugar this is expressed as:

$$M_{sugar,i} = c_{sugar,i} \times M_i = V_i \times \rho_w \times c_{sugar,i}$$

The mass of a hollow cylinder representing the outer most centimetre of xylem biomass (e.g., 0-1 cm; $M_{i,0}$) is derived from its volume ($V_{i,0}$) and an average wood density ($\rho_w$). Here, $\rho_w = 289.9 \pm 40.4\ kg\,m^{-3}$ was used as wood density, which is the mean ± standard deviation of wood density of 16 cores from nine trees in the control group as determined using x-ray film according to (Björklund et al., 2019). The volume is calculated using a height ($h$) of 10 cm and the section's circumference ($cbh_i$; in $cm$), which was measured in the field using a tape measure.

$$M_{i,0} = V_{i,0} \times \rho_w = \pi \times h \times \rho_w \Big( \Big(\frac{cbh_i}{2\pi}\Big)^2 - \Big(\frac{cbh_i}{2\pi} - 0.01\Big)^2 \Big)$$

Analogously, we calculated the mass of a hollow cylinder representing the second outer most centimetre of material (e.g., 1-2 cm; $M_{i,1}$) as follows.

$$M_{i,1} = V_{i,1} \times \rho_w = \pi \times h \times \rho_w \left( \left( \frac{cbh_i}{2\pi} - 0.01 \right)^2 - \left( \frac{cbh_i}{2\pi} - 0.02 \right)^2 \right)$$

For example, a 10 $cm$ high stem section of 20 $cm$ diameter, would have a volume of 628 $cm^3$ and a mass of 182.06 $g$. At an average measured sugar and starch concentration of 0.83 and 0.28 $\% \, dry \, mass$ in July, this equates to a total nonstructural carbon reserve of roughly 2 $g$ or 1 $g$ of easily accessible carbon in the first two centimetres of that stem section.
